# Distribution, organization and expression of genes concerned with anaerobic lactate-utilization in human intestinal bacteria

**DOI:** 10.1101/2021.04.04.438253

**Authors:** Paul O. Sheridan, Petra Louis, Eleni Tsompanidou, Sophie Shaw, Hermie J. Harmsen, Sylvia H. Duncan, Harry J. Flint, Alan W. Walker

## Abstract

Lactate accumulation in the human gut is linked to a range of deleterious health impacts. However, lactate is consumed and converted to the beneficial short chain fatty acids butyrate and propionate by indigenous lactate-utilizing bacteria. To better understand the underlying genetic basis for lactate utilization, transcriptomic analysis was performed for two prominent lactate-utilizing species from the human gut, *Anaerobutyricum soehngenii* and *Coprococcus catus*, during growth on lactate, hexose sugar, or hexose plus lactate. In *A. soehngenii* L2-7, six genes of the *lct* cluster including NAD-independent D-lactate dehydrogenase (i-LDH) were co-ordinately upregulated during growth on equimolar D and L-lactate (DL-lactate). Upregulated genes included an acyl-CoA dehydrogenase related to butyryl-CoA dehydrogenase, which may play a role in transferring reducing equivalents between reduction of crotonyl-CoA and oxidation of lactate. Genes upregulated in *C. catus* GD/7 included a six-gene cluster (*lap*) encoding propionyl CoA-transferase, a putative lactoyl-CoA epimerase, lactoyl-CoA dehydratase and lactate permease, and two unlinked acyl-CoA dehydrogenase genes that are candidates for acryloyl-CoA reductase. An i-LDH homolog in *C. catus* is encoded by a separate, partial *lct,* gene cluster, but not upregulated on lactate. While *C. catus* converts three mols of DL-lactate via the acrylate pathway to two mols propionate and one mol acetate, some of the acetate can be re-used with additional lactate to produce butyrate. A key regulatory difference is that while glucose partially repressed *lct* cluster expression in *A. soehngenii*, there was no repression of lactate utilization genes by fructose in the non-glucose utilizer *C. catus.* This implies that bacteria such as *C. catus* might be more important in curtailing lactate accumulation in the gut.

**Impact statement:** Lactate can be produced as a fermentation by-product by many gut bacteria but has the potential to perturb intestinal microbial communities by lowering luminal pH, and its accumulation has been linked to a range of deleterious health outcomes. Fortunately, in healthy individuals, lactate tends not to accumulate as it is consumed by cross-feeding lactate-utilizing bacteria, which can convert it into the beneficial short chain fatty acids butyrate and propionate. Lactate-utilizing gut bacteria are therefore promising candidates for potential development as novel probiotics. However, lactate-utilizers are taxonomically diverse, and the genes that underpin utilization of lactate by these specialized gut bacteria are not fully understood. In this study we used transcriptomics to compare gene expression profiles of *Anaerobutyricum soehngenii* and *Coprococcus catus,* two prominent lactate-utilizing species in the human gut, during growth on lactate alone, sugar alone, or sugar plus lactate. The results revealed strong upregulation of key, but distinct, gene clusters that appear to be responsible for lactate utilization by these, and other, gut bacterial species. Our results therefore increase mechanistic understanding of different lactate utilization pathways used by gut bacteria, which may help to inform selection of optimal lactate-utilizing species for development as novel therapeutics against colonic microbiota perturbations.

**Data summary:** Novel draft genomes generated for this study have been made available from GenBank (https://www.ncbi.nlm.nih.gov/bioproject/) under BioProject number PRJNA701799. RNA-seq data have been deposited in the ArrayExpress database at EMBL-EBI (www.ebi.ac.uk/arrayexpress) under accession number E-MTAB-10136. Further details of additional existing genomic data that were analyzed in this project are given in Table 1 and Table S2.

## Introduction

The human large intestinal microbiota is dominated by obligately anaerobic bacteria, whose growth is largely dependent on the supply of complex carbohydrates and proteins available for fermentation. These substrates are mostly fermented by resident gut anaerobes to short chain fatty acids (SCFAs) and gases, and in the healthy colon the major faecal SCFAs detected are acetate, propionate and butyrate [1]. Lactate is another fermentation product of many anaerobic bacteria that colonize the mammalian gut, either as a major metabolite, as in *Lactobacillus* and *Bifidobacterium* spp., or as an alternative hydrogen sink [2]. However, as the pKa (3.73) for lactic acid is lower than that of other fermentation acids such as acetate (4.76), accumulation of lactate has the potential to dramatically change the gut environment and gut microbiota composition via reduction in prevailing pH [3].

Lactate may therefore help to inhibit the growth of some pathogenic bacteria with poor tolerance for lower pHs [4] and a gut microbiota dominated by lactate-producing bacteria is found in many healthy pre-weaned infants [5]. On the other hand, there are many known deleterious impacts of lactate accumulation in the gut. Indeed, the phenomenon of lactic acidosis, driven by excessive production of lactate following fermentation of easily digestible dietary carbohydrates, is well known in ruminants and is a major problem in animal husbandry [6, 7]. Similarly, in humans, accumulation of toxic D-lactate is life-threatening in short bowel syndrome [8]) and lactate accumulation is also associated with severe colitis [9]. Additionally, lactate can be utilized as a carbon and energy source by certain intestinal pathogens such as *Salmonella* [10] and *Campylobacter* [11], compounding the potential heath detriments of lactate accumulation in the gut.

However, in the colon of healthy adult humans, as well as in the non-acidotic rumen, lactate does not accumulate, due the activities of cross-feeding, lactate-utilizing bacteria [12]. Colonic lactate concentrations are therefore a balance between bacterial production during fermentation and utilization by lactate-utilizing bacteria, which can use lactate to form acetate, propionate or butyrate [13]. Recent research has linked the populations and activities of these lactate-utilizing bacteria with overall fermentation patterns and productivity [14], and a low abundance of commensal lactate-utilizers has also been proposed to make gut microbial communities inherently less stable and more prone to lactate-induced perturbations [15]. As such, lactate-utilizing bacteria have been proposed as promising candidates for development as novel probiotics [15–18].

The ability to utilize lactate as an energy source for growth appears to be limited to relatively few bacterial species among the human intestinal microbiota, although these species are taxonomically diverse and can utilize lactate in different ways. Selective isolation on DL lactate-containing media resulted in the recovery of *Lachnospiraceae* species subsequently identified as *Eubacterium hallii* (since reclassified as the two species *Anaerobutyricum hallii* and *Anaerobutyricum soehngenii* [19]), *Anaerostipes hadrus* and *Anaerostipes caccae* [20, 21]. These isolates produce butyrate from lactate, with net consumption of acetate, suggesting that they initially convert lactate into pyruvate, which is then routed into the butyrate pathway [21]. The lactate to pyruvate conversion is energetically unfavourable and recent evidence in *Acetobacterium woodii* indicates that it is dependent on electron confurcation [22]. Alternative routes for lactate utilization that are known to be important in the rumen [23] result in propionate formation. These include the acrylate pathway found in the *Negativicutes* species *Megasphaera elsdenii* and in at least one member of the *Lachnospiraceae*, *Coprococcus catus* [24], and the succinate pathway that is found among the *Negativicutes* in *Selenomonas ruminantium* and *Veillonella* spp. [24]. Less advantageously, lactate is also a co-substrate for sulfate-reducing bacteria that use it to form acetate and sulfide, the latter of which may be genotoxic [25].

Here we examine the distribution, organization and regulation of genes involved in lactate utilization among dominant representatives of the human intestinal microbiota, based on new isolates of lactate-utilizing species and on newly available genome sequences. In particular, we identify two gene clusters whose transcription is upregulated during growth with lactate as carbon and energy source. One of these corresponds to the *lct* gene cluster of *A. woodii* [22] that was recently identified from proteomic analysis in *A. soehngenii* [26] while the second encodes activities involved in the acrylate pathway of *Coprococcus catus*. This investigation reveals differences in the regulation, phylogenetic distribution and metabolic function of these two clusters, and provides novel insights into the mechanistic basis of different lactate-utilization strategies used by human gut bacteria.

## Methods

### Isolation of new strains of lactate utilizing bacteria

New strains of *Anaerobutyricum soehngenii* (HTF-83D) and *Anaerostipes hadrus* (HTF-920, HTF-146, HTF-370 and HTF-412) were isolated from human faecal samples by dilution and culturing of single colonies on the anaerobic medium YCFA [27] supplemented with glucose. Taxonomic identification of strains was carried out by 16S rRNA gene sequencing and BLASTn querying each sequence against the NCBI 16S rRNA gene reference database. New strains of *Anaerobutyricum* were further classified as either *A. hallii* or *A. soehngenii* by average nucleotide identity (ANI) and average amino acid identity (AAI) using nucmer [28] and CompareM (https://github.com/dparks1134/CompareM), respectively (Table S1).

### Bacterial strains, growth conditions and genomes

The bacterial strains and genomes used in this work, including both new and previously isolated strains, are described in Table 1 and Table S2. Routine culturing of bacterial strains was in anaerobic M2GSC medium [29] in 7.5 ml aliquots in Hungate tubes, sealed with butyl rubber septa (Bellco Glass). Growth experiments were carried out in basal YCFA medium [27] supplemented with 35 mM lactate and/or 11 mM glucose (or fructose for *C. catus* since this species is incapable of growing on glucose as sole carbon source). All cultures were incubated anaerobically without agitation at 37°C, using the anaerobic methods described previously by Bryant [30].

**Table 1.**
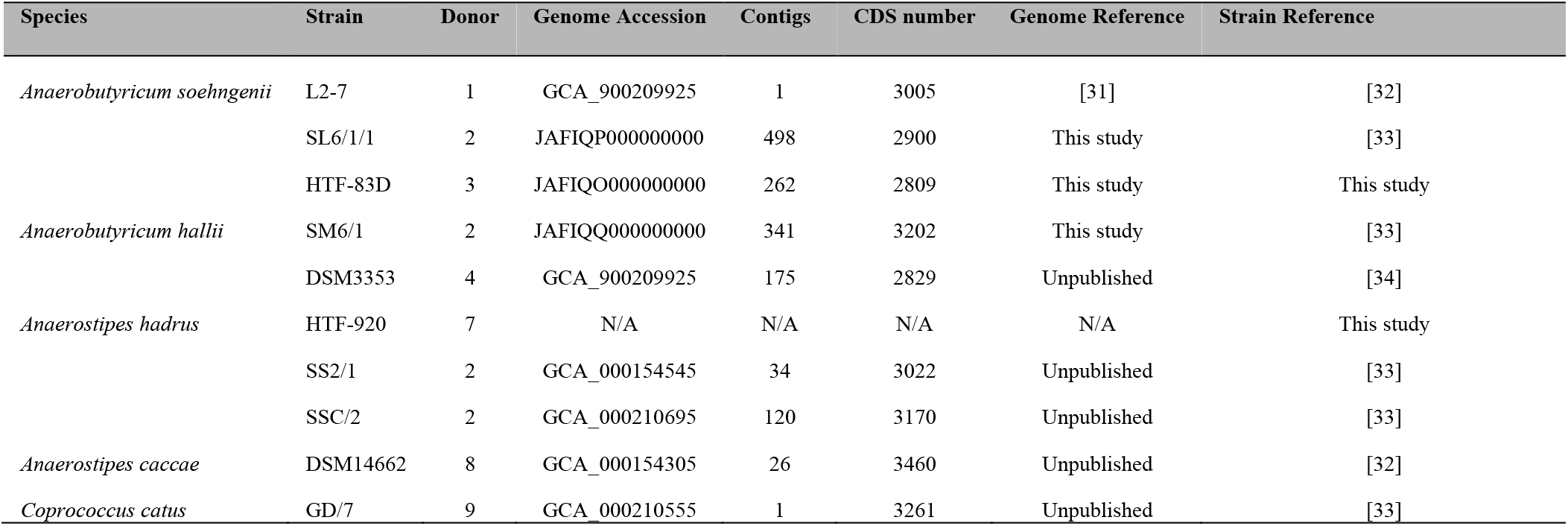
New and existing strains used for in vitro cultivation studies in this work, and their corresponding genomes. “N/A” indicates no genome available for this strain, which was used for *in vitro* work only. The full list of strains used for genomic-based analyses is shown in Table S2.

### DNA isolation, genome sequencing and annotation

Genomic DNA was extracted with the Ultraclean Microbial DNA isolation kit (MoBio Laboratories, Carlsbad, CA, USA). The DNA concentration and purity were measured using a NanoDrop 2000c spectrophotometer (Thermo Fisher Scientific, Waltham, MA, USA) and the Qubit double-stranded DNA (dsDNA) HS and BR assay kits (Life Technologies, Carlsbad, CA, USA). One nanogram of bacterial DNA was used for library preparation. The DNA library was prepared using the Nextera XT library preparation kit with the Nextera XT v2 index kit (Illumina, San Diego, CA, USA). The library fragment length was aimed at fragments with a median size of 575 bases and was assessed with the Genomic DNA ScreenTape assay with the 2200 Tape-Station system (Agilent Technologies, Waldbronn, Germany). Subsequently, the library was sequenced on an Illumina MiSeq sequencer, using a 2×250 (500v2) cartridge, using the MiSeq reagent kit v2 generating 250-bp paired-end reads, (Illumina, San Diego, CA, USA). MiSeq data were processed with MiSeq control software v2.4.0.4 and MiSeq Reporter v2.4 (Illumina, San Diego, CA, USA). The resulting fastq files were filtered using TrimGalore v0.4.0 (https://github.com/FelixKrueger/TrimGalore), removing one nucleotide off the 3’ end (--trim1) and removing both pairs of the paired end reads if one read did not pass filtering (--paired). Filtered fastq files were assembled using the idba_ud assembler v1.1.3 [35] using standard settings. Newly generated genome sequences are available from GenBank (BioProject PRJNA701799), with individual accession numbers shown in Table 1 and Table S2. Additional sequences were retrieved from GenBank (accession numbers in Table S2). New genome sequences were annotated using Prokka v1.11 [36] and publicly available sequences were reannotated in the same manner to ensure uniformity between annotation methods. Genome completeness was estimated using CheckM v1.1.3 [37].

### Genome analyses

Lactate utilization loci were searched for using the Hidden Markov Model (HMM) profiles of conserved domains found in three-component L-lactate hydrogenase (PF02754, PF02589, PF13183, PF11870), FAD-dependent D-lactate dehydrogenase (PF01565, PF02913, PF09330, PF12838, PF02754), FMN-dependent L-lactate dehydrogenase (PF01070, PF00173), NAD-dependent D-lactate dehydrogenase (PF00389, PF02826), NAD-dependent L-lactate dehydrogenase (PF02866, PF00056), lactate permease (PF02652), lactate racemase (PF09861) and ETF (PF00766, PF01012) using hmmsearch in HMMER3 [38]. Butyrate and propionate producing pathways were detected by BLASTP against UniProt [39] and KEGG [40] with a threshold of 60% identity using the marker genes butyrate kinase and butyryl-CoA:acetate CoA-transferase as indicators of butyrate production, and propanediol utilization protein (PduP), lactoyl CoA dehydratase alpha-subunit (LcdA) and methyl-malonyl-CoA decarboxylase alpha-subunit (MmdA) as indicators of propionate production through the propanediol, acrylate and succinate pathways, respectively [24]. Rnf complexes were detected by query against amino acid sequences of the experimentally proven Rnf complexes of *Clostridium ljungdahlii* and *A. woodii* [41, 42].

All genes containing the lactate permease Pfam [43] domain PF02652 in UniProt (June 2020) [39] were downloaded and clustered into groups of >70% similarity using CD-HIT v4.8.1 [44]. Representatives of each cluster were aligned using MAFFT L-INS-I v7.407 [45]. The acyl-CoA dehydrogenase in the *lct* cluster (L2-7_01909) was used as a BLASTP query against *C. catus* GD/7 and the *A. hadrus* genomes, homologs were aligned using MAFFT L-INS-i [45] and a HMM profile was created from the alignment using hmmbuild in HMMER3 [38]. This profile was then queried against the genomes of known lactate utilizers using hmmsearch (-T 80) to detect divergent members of the acyl-CoA dehydrogenase protein superfamily. Matching sequences were aligned using MAFFT L-INS-i [45] and the alignment was manually inspected for likely mis-annotations (none were found). For both the lactate permease and acyl-CoA dehydrogenase alignments, spurious sequences and poorly aligned regions were removed with trimal (automated1, resoverlap 0.55 and seqoverlap 60) [46] and maximum likelihood trees were constructed with IQ-TREE [47] using the best fitting protein model predicted in ModelFinder [48]. Branch supports were computed using the SH-aLRT test [49] for the lactate permease tree and ultrafast bootstraps for the acyl-CoA dehydrogenase tree. The tree figures were generated using iTOL [50].

### Differential gene expression analysis

Fresh YCFA broth supplemented with 11 mM monosaccharide (glucose for *A. soehngenii* and fructose for *C. catus*), 35 mM lactate or both 11 mM monosaccharide and 35 mM lactate were inoculated 1:75 with an overnight grown culture of either *A. soehngenii* L2-7 or *C. catus* GD/7. These cultures were grown to OD650 0.3 before cultures were centrifuged to cell pellets (5 min, 1,200 *g*, 20 °C). The supernatant was transferred to a new tube for SCFA and monosaccharide analysis, and the cell pellet was resuspended in 500 µl of RNAlater (Invitrogen™), left at room temperature for 5 mins and frozen at −70 °C. Total RNA was extracted using the RNeasy PowerMicrobiome Kit (QIAGEN), following the manufacturer’s instructions. This included a bead beating step for lysing Gram-positive cell walls. Ribosomal RNA was depleted using the Illumina Ribo-Zero rRNA removal kit. Libraries were prepared using the Illumina TruSeq stranded mRNA library kit and sequenced at the Centre for Genome-Enabled Biology and Medicine (CGEBM) at The University of Aberdeen using a High Output 1×75 kit on the Illumina NextSeq platform producing 75 bp single end reads. Reads were combined from two runs producing between 14237482 and 16008520 reads per sample. These transcriptomic datasets are available from the ArrayExpress database at EMBL-EBI under accession number E-MTAB-10136. Quality of raw reads was assessed using FastQC v0.11.3 [51] and sequences were filtered using TrimGalore v0.4.0. Between 14141677 and 15921037 sequences remained per sample, which, on average, was 99% of the raw reads. The appropriate reference genomes for *A. soehngenii* L2-7 and *C. catus* GD/7 (Table S2) were indexed in preparation for alignment using the HISAT2 build function (version 2.1.0) [52]. Filtered reads were then aligned to the appropriate reference genome using HISAT2 v2.1.0 [52] with default settings. Alignments were then converted to BAM format and sorted using SAMtools v1.2 [53]. Reads aligned to gene regions were counted using featureCounts (subread package version 5.0-pl) [54] with settings to split multi-mapping reads as a fraction between aligned regions. Genes with significant differential expression (FDR < 0.05) between conditions were identified using edgeR v3.16.5 [55], utilizing the GLM function, as more than two growth conditions were used in this study.

### Short chain fatty acid, monosaccharide and ethanol analysis

SCFA concentrations were measured by gas chromatography (GC) as described previously (Richardson *et al*., 1989). In brief, following derivatization of the samples using N-tert-butyldimethylsilyl-N-methyltrifluoroacetamide, the samples were analysed using a Hewlett Packard gas chromatograph (GC) fitted with a silica capillary column using helium as the carrier gas. L-lactate concentration was assessed by Lactate Reagent (Trinity biotech) using a Konelab 30 chemistry analyser. D-lactate was calculated as the difference between total lactate and L-lactate. Methanol, ethanol, propanol, butanol and pentanol concentrations were also measured by GC using a ZB WAX column. Glucose and fructose concentration were assessed using the glucose hexokinase and fructose hexokinase assays for Konelab.

## Results

### Lactate utilization in human colonic bacteria

Some anaerobic colonic bacteria are capable of utilizing lactate as a carbon and energy source. This process can generate the SCFAs butyrate, propionate or acetate depending on the species [21, 24]. Previous studies have been based on only a small number of available strains, but the isolation and genome sequencing here of new strains of lactate-utilizing species allowed for more in-depth analysis. Each strain was grown in culture provided with glucose (or fructose in the case of *C. catus* GD/7), DL-lactate, or a combination of the two as energy sources (Figure 1). All nine strains identified as either *Anaerostipes* or *Anaerobutyricum* spp. showed the ability to consume lactate and acetate with production mainly of butyrate, as previously reported [21]. This contrasts with production mainly of propionate from lactate by *C. catus* (Figure 1). The *A. hadrus* strains tested showed little or no ability to utilize L-lactate, whereas the other species utilized both isomers (Figure 1B). Lactate-utilization was repressed by the presence of hexose sugar in the growth medium to varying extents for the different *Anaerobutyricum* and *A. hadrus* strains, with utilization of both lactate stereoisomers particularly repressed by the presence of glucose for *A. soehngenii* strains L2-7 and SL6/1/1 (Figure 1B). Interestingly, although propionate was the main SCFA produced by *C. catus* GD/7 when grown on lactate some butyrate and acetate production was also observed, indicating the presence of these two SCFAs as intermediate- or end-products of lactate utilization in this organism.

**Figure 1.**
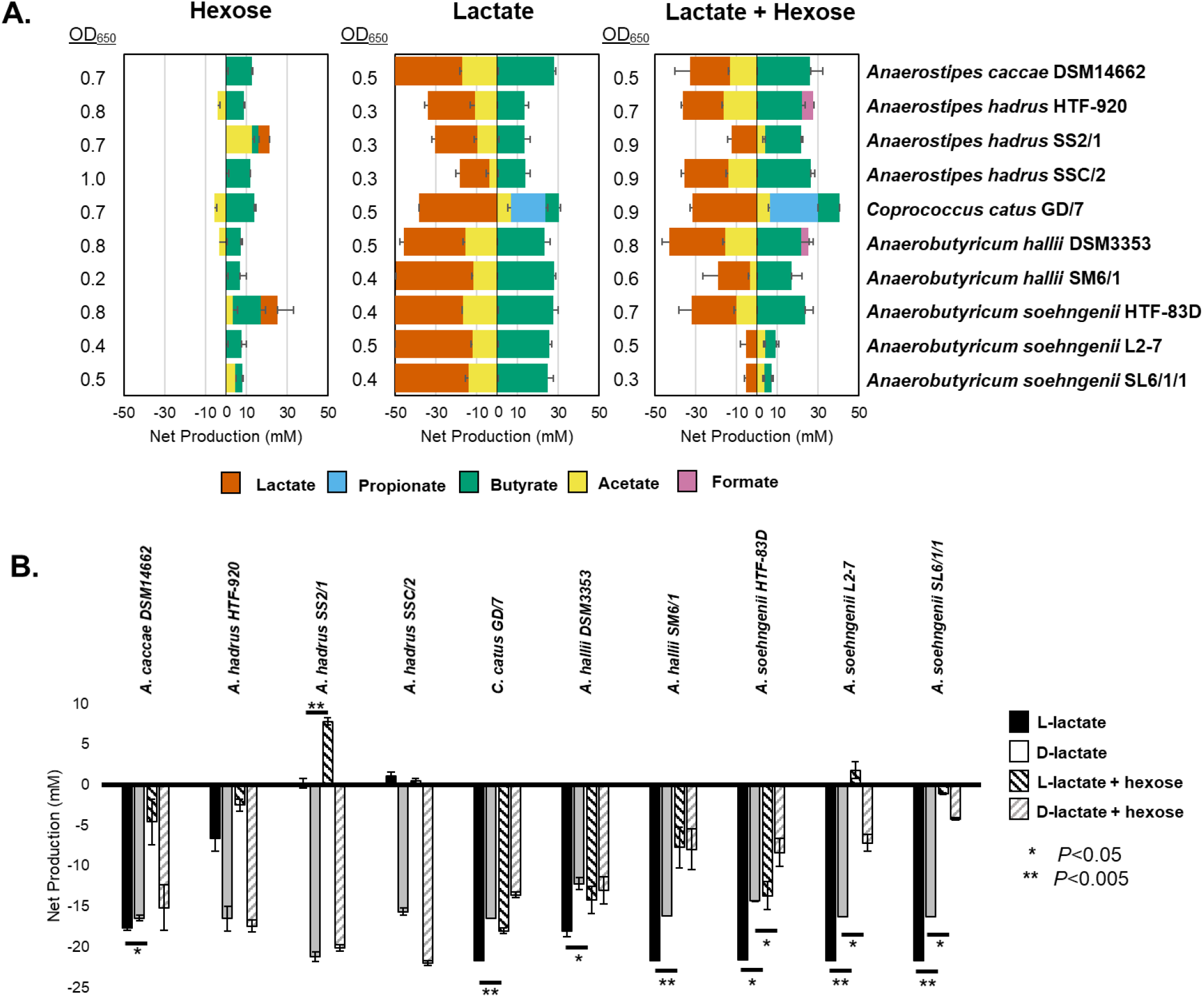
Fermentation profiles of lactate-utilizing bacteria grown on sugar, on lactate, or a combination of both. Panel A shows the net production (to the right of the figures), or consumption (to the left of the figures) of various short chain fatty acids by lactate-utilizing bacteria grown on either hexose sugar alone, lactate alone, or a combination of both. Cultures were incubated anaerobically for 24 h in media supplemented with 11 mM glucose, 35 mM lactate or both. Glucose was substituted with 11 mM fructose for *C. catus* GD/7. Iso-butyrate, valerate, iso-valerate, and succinate were not detected in any of the samples. Panel B shows net consumption or production of L- or D-lactate by the tested strains, with or without simultaneous exposure to hexose sugars in the culture medium. Error bars are the SEM of three biological replicates. There was no remaining glucose or fructose in any culture, with the exception of *A. soehngenii* L2-7 (2 mM, SEM 2.0) and SL6/1/1 (6 mM, SEM 3.2). Expanded data are shown in Table S3.

### Changes in gene expression during growth on DL-lactate

To investigate the genetic control and regulation of lactate utilization, transcriptomic analysis was performed on two prevalent lactate-utilizing species from the human gut, *A. soehngenii* and *C. catus*. This involved assessing the differential expression of genes during growth on a hexose sugar (glucose for *A. soehngenii* and fructose for *C. catus*), on DL-lactate or on a combination of sugar plus DL-lactate (Figure S1). Growth was carefully monitored in each of six replicate experiments for each strain to ensure sampling during mid-exponential growth phase (OD650=0.3). RNA was extracted from cell pellets and the culture supernatant was analysed for their SCFA profiles and remaining hexose (Figure 2). In agreement with the results shown in Figure 1, in the absence of glucose *A. soehngenii* L2-7 had utilized much of the lactate at the time of sampling, but lactate utilization was significantly repressed in the presence of glucose (*P* < 0.001). In the absence of hexose sugar *C. catus* GD/7 had also utilized much of the lactate, but utilization was slightly lower in the presence of fructose (*P* < 0.01). Sugar was still present in the growth media initially supplemented with it (Figure 2), indicating there was potential for repression at the point of RNA harvesting.

**Figure 2.**
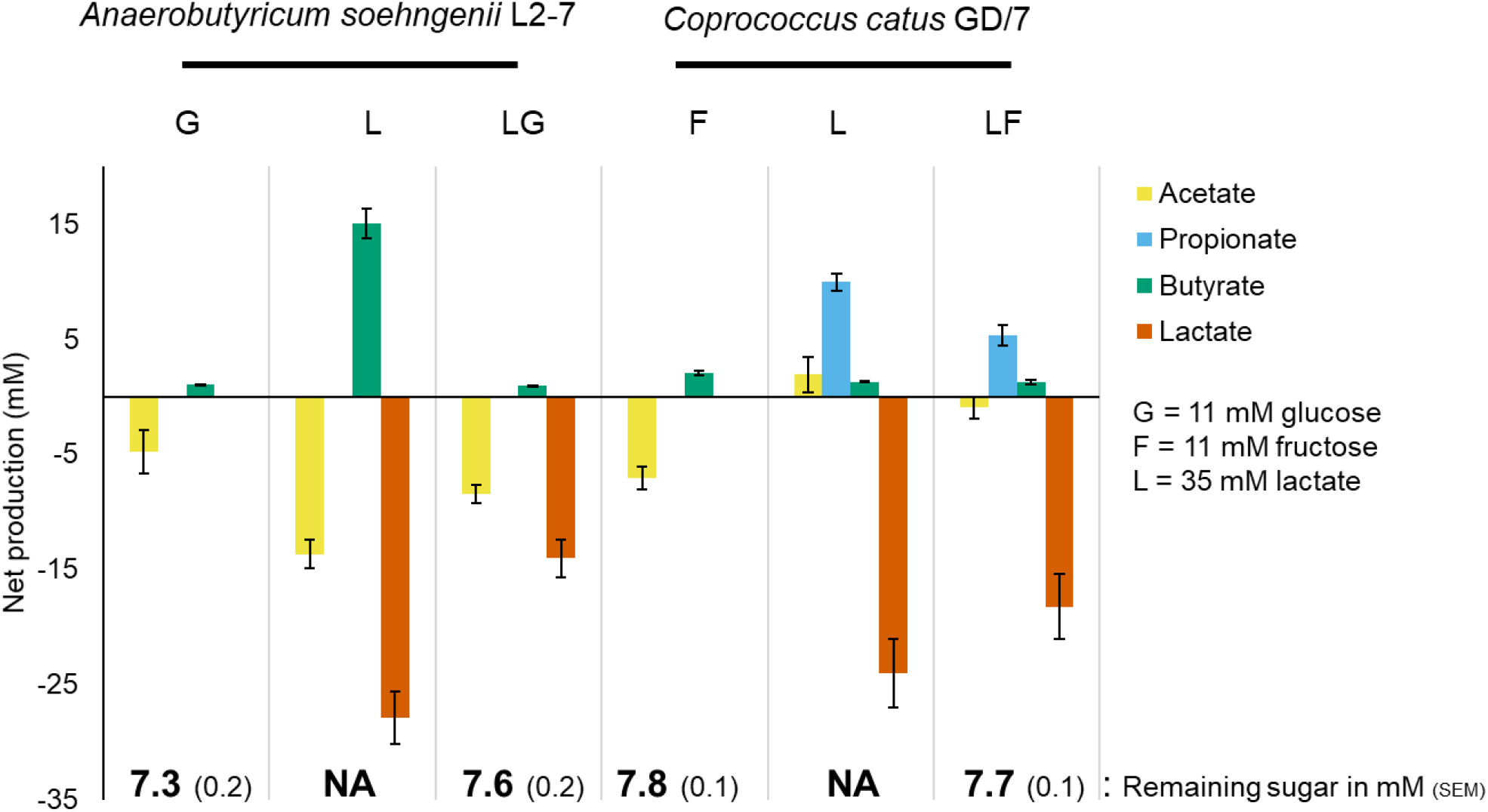
Fermentation end products of *A. soehngenii* L2-7 and *C. catus* GD/7 during growth on lactate, a monosaccharide alone or on both in combination. Cultures were harvested at an OD650 reading of 0.3. Iso-butyrate, valerate, iso-valerate and succinate were not detected in any of the samples. Error bars are the SEM of six biological replicates.

The results from the experiments with *A. soehngenii* L2-7 showed differences in overall transcriptomic profile were observed during growth on lactate compared to glucose alone or a combination of glucose and lactate (Figure 3). Conversely, little overall difference in transcriptomic profile was observed between glucose and the combination of glucose and lactate, with only 24 differentially expressed genes as opposed to 1427 between glucose and lactate as a sole carbon source. In *C. catus* GD/7, the overall transcriptomic profile was again strongly influenced by the growth substrate (Figure 3). In *C. catus* GD/7, however, and in contrast to *A. soehngenii* L2-7, lactate-induced genes were not simply repressed by the presence of monosaccharide (fructose), with all three transcriptome profiles differing from each other.

**Figure 3.**
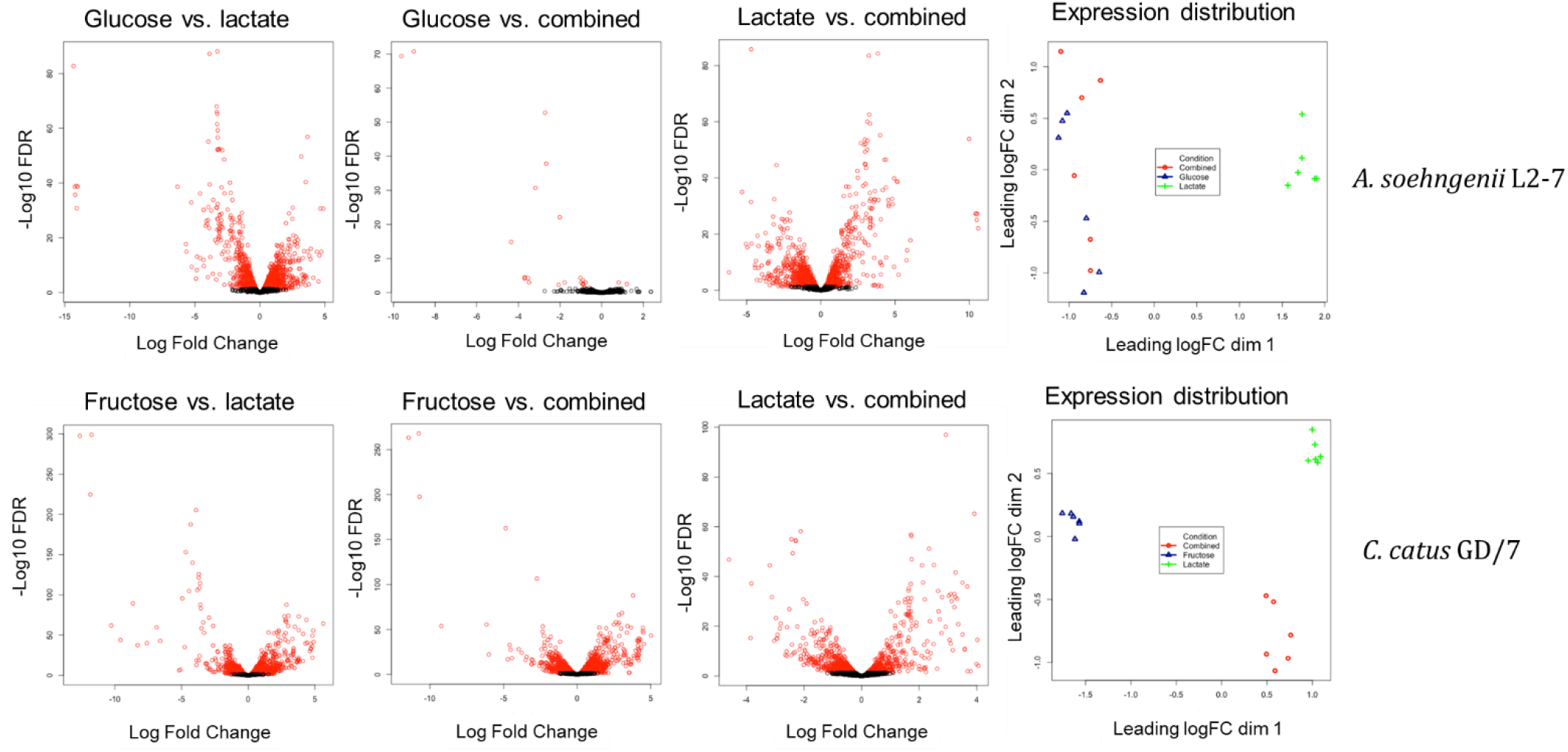
Volcano plots of log fold gene expression change vs. –log10 of the false discovery rate value and multidimensional scaling plots showing the gene expression distribution of samples during growth on lactate, monosaccharide or a combination of the two. Genes with a significant FDR value <0.05 in the volcano plots are highlighted in red. A more detailed overview is given in Table S4. “Combined” indicates cultures incubated with both lactate and hexose.

In *A. soehngenii* L2-7, eleven transcripts showed >5 Log2-fold increase in expression during growth on lactate alone compared with growth on glucose alone, and increases were lower when glucose was present alongside lactate (Table 2). Six transcripts (L2-7_01905-01910) that showed the highest coordinate upregulation (log2 >14) were encoded by the *lct* gene cluster that includes an NAD-independent lactate dehydrogenase (iLDH), lactate permease, ETF electron transport factor (alpha and beta subunits), lactate racemase and an acyl-CoA dehydrogenase (Table 2). The linked regulatory gene *lctA* showed a lower amplitude of induced expression. A second cluster of three genes that showed the same behaviour, also at lower amplitude, included an autoinducer, RNA polymerase sigma factor and HTH regulatory protein. Two further transcripts, from a second lactate racemase and closely linked aquaporin genes, were also highly induced by lactate, but were not repressed by glucose. The components of the Rnf complex, which couples the oxidation of reduced ferredoxin to the reduction of NAD [41], was highly induced by lactate and repressed by glucose, as were the components of the Hnd complex, which catalyzes the reduction of NADP in the presence of molecular H2 to yield NADPH [56, 57].

**Table 2.** Selected genes with high expression differences during growth on glucose (G), fructose (F), lactate (L) or a combination of substrates (LG/LF). All transcripts are listed that exhibited >5 log2-fold changes in expression, together with expression changes for linked genes within the same gene cluster. Changes of less than 5-fold are listed for the Rnf and Hnd genes clusters in *A soehngenii* because of their metabolic relevance. All listed genes have an FDR value of <0.05. NoDiff = No significant difference in expression. Expanded to all genes in Tables S5-S10.

| *A. soehngeni* L2-7 |  | log2-fold change |  |  |
| --- | --- | --- | --- | --- |
|  |  | G v L | G v LG | L v LG |
| L2-7_01905 | Lactate permease (*lctE*) | -14.3 | -4.4 | 10.0 |
| L2-7_01906 | D-iLDH (*lctD*) | -14.2 | -3.7 | 10.5 |
| L2-7_01907 | ETF-B (*lctB*) | -14.1 | -3.5 | 10.6 |
| L2-7_01908 | ETF-A (*lctC*) | -14.2 | -3.7 | 10.5 |
| L2-7_01909 | Acyl-CoA dehydrogenase (*lctG*) | -14.1 | -3.7 | 10.4 |
| L2-7_01910 | Lactate racemase (*lctF*) | -14.0 | -3.5 | 10.5 |
| L2-7_01911 | LutR transcriptional regulator (*lctA*) | -3.0 | NoDiff | 2.5 |
| L2-7_02157 | Aquaporin family protein | -6.3 | -9.6 | -3.3 |
| L2-7_02158 | Racemase | -5.3 | -9.0 | -3.7 |
| L2-7_02594 | Putative cyclic lactone autoinducer peptide | -5.2 | NoDiff | 5.8 |
| L2-7_02595 | Hypothetical protein | -4.9 | NoDiff | 4.9 |
| L2-7_02596 | RNA polymerase subunit sigma-70 | -5.7 | NoDiff | 6.0 |
| L2-7_02597 | Helix-turn-helix domain-containing protein | -5.7 | NoDiff | 5.8 |
| L2-7_00821 | Hnd complex, subunit A | -4.33 | NoDiff | 5.12 |
| L2-7_00822 | Hnd complex, subunit C | -4.07 | NoDiff | 4.56 |
| L2-7_00823 | Hnd complex, subunit D | -3.98 | NoDiff | 4.38 |
| L2-7_00377 | Rnf complex, subunit C | -2.90 | NoDiff | 2.91 |
| L2-7_00378 | Rnf complex, subunit D | -3.08 | NoDiff | 2.98 |
| L2-7_00379 | Rnf complex, subunit G | -3.20 | NoDiff | 3.13 |
| L2-7_00380 | Rnf complex, subunit E | -3.33 | NoDiff | 3.25 |
| L2-7_00381 | Rnf complex, subunit A | -3.27 | NoDiff | 3.22 |
| L2-7_00382 | Rnf complex, subunit B | -3.24 | NoDiff | 3.15 |

**Table 2. Selected genes with high expression differences during growth on glucose (G), fructose (F), lactate (L) or a combination of substrates (LG/LF).**
| *C. catus* GD/7 |  | log2-fold change |  |  |
| --- | --- | --- | --- | --- |
|  |  | F v L | F v LF | L v LF |
| GD-7_01094 | Propionyl CoA transferase (*lapA*) | -10.3 | -9.2 | 1 |
| GD-7_01095 | Lactoyl CoA epimerase (*lapB*) | -11.8 | -10.7 | 1.1 |
| GD-7_01096 | Lactoyl CoA dehydratase subunit (*lapC*) | -12.6 | -11.4 | 1.2 |
| GD-7_01097 | Lactoyl CoA dehydratase subunit (*lapD*) | -13.1 | -11.9 | 1.2 |
| GD-7_01098 | Lactoyl CoA dehydratase subunit (*lapE*) | -13 | -11.9 | 1.2 |
| GD-7_01099 | Lactate permease (*lapF*) | -13 | -11.9 | 1.2 |
| GD-7_00065 | 3-hydroxybutyryl-CoA dehydratase | -8.6 | -6.2 | 2.5 |
| GD-7_00066 | H<sup>+</sup>/gluconate symporter and related permeases | -6.9 | -4.6 | 2.3 |
| GD-7_00760 | Amidohydrolase family protein | -7.6 | -4.4 | 3.2 |
| GD-7_00761 | MFS transporter | -8.3 | -4.7 | 3.6 |
| GD-7_01087 | Acyl-CoA dehydrogenase | -9.6 | -6 | 3.6 |
| GD-7_01088 | ABC transporter substrate-binding protein | -6.6 | -3.5 | 3.1 |
| GD-7_01551 | Acyl-CoA dehydrogenase | -11.7 | -10.8 | 1 |

In *C. catus* GD/7, seven genes showed >10 Log2 fold increased expression during growth on lactate alone compared with fructose alone (Table 2). Six of these were encoded by a cluster of linked genes, but this cluster showed no sequence similarity with the *lct* cluster of *A. soehngenii* described above and comprised genes (GD-7_01094-01099) encoding three components of lactoyl-CoA dehydratase, propionyl-CoA transferase, a methyl-malonyl-CoA epimerase homolog and a lactate permease (Table 2). These genes encode most of the activities required for the acrylate pathway to convert lactate into propionate, consistent with previous evidence using isotopically labelled lactate showing that this bacterium uses the acrylate pathway for lactate utilization [24]. This locus is thus a cluster encoding lactate utilization via the acrylate pathway (*lap*). It seems likely that the methyl-malonyl-CoA epimerase homolog (*lapB*) functions as a lactoyl-CoA epimerase, catalyzing the reversible conversion of L-lactoyl-CoA to D-lactoyl-CoA. The upregulated lactate permease gene was one of four encoded at separate locations in the *C. catus* GD/7 genome (Table S11). These six genes, together with two unlinked genes (GD-7_01551, GD-7_01087) encoding acyl-CoA dehydrogenases, were also highly upregulated on lactate when fructose was present. The two acyl-CoA dehydrogenases are candidates for propionyl-CoA dehydrogenase (acrylate reductase) activity.

### Wider occurrence of lactate-utilization gene clusters in the genomes of human colonic bacteria

Homologs of the putative lactate utilization genes identified in *A. soehngenii* L2-7 and *C. catus* GD/7 were also detected in other known lactate-utilizing bacteria from the human colonic microbiota and other environments (Figure 4A & B, Table S12 & S13). Homologs of the upregulated *lct* genes from *A. soehngenii* L2-7 were present in the two species of *Anaerobutyricum*, *Anaerostipes caccae*, *Eubacterium limosum*, *C. catus* and *Megasphaera elsdenii* (Table S12) (although as noted above they were upregulated only moderately in *C. catus* (Table S8)). The genome for *A. hadrus* DSM3319 lacked the racemase gene, consistent with the observation that it consumed only the D isomer of lactate (Figure 1B). *Lct* homologs were also present in the *Acetobacterium woodii* gene cluster, but this cluster does not contain the putative acyl-CoA dehydrogenase (ACoAD) gene *lctG*.

**Figure 4.**
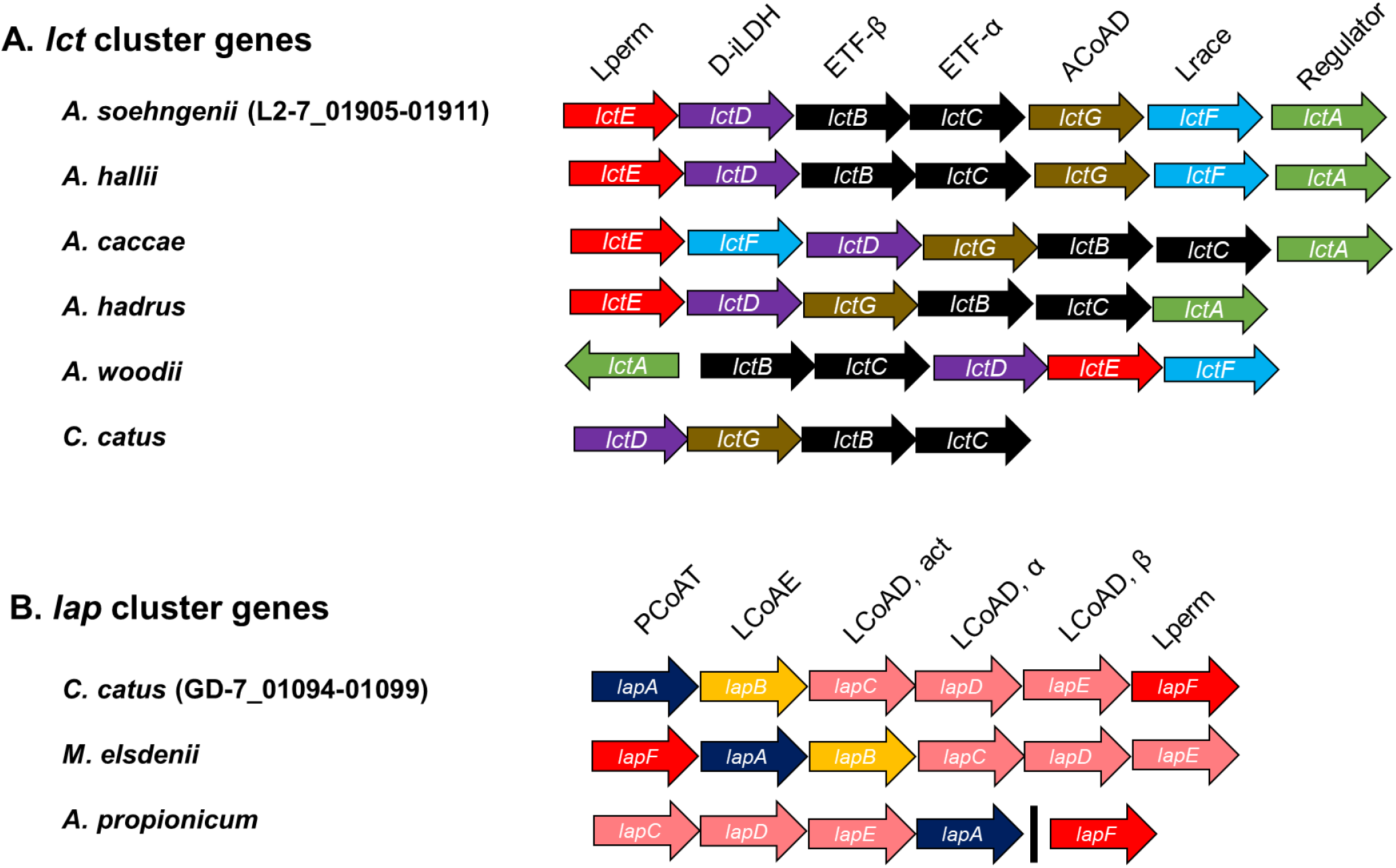
Organization of lactate utilization loci observed in (A) *A. soehngenii* and (B) *C. catus* in other previously described lactate-utilizing bacteria. The black line in the *A. propionicum lap* cluster indicates that *lapF* gene is present in a different region of the genome. Lactate permease (Lperm), NAD-independent D-lactate dehydrogenase (D-iLDH), electron transfer flavoprotein (ETFα & β), acyl-CoA dehydrogenase (ACoAD), lactate racemase (Lrace), lactate operon transcriptional regulator (Regulator) propionyl-CoA transferase (PCoAT), lactoyl CoA epimerase (LCoAE), lactoyl-CoA transferase (LCoAD, act, α & β). Expanded to more lactate utilizing bacteria genomes in Tables S12-13.

Homologs of the six clustered *lap* cluster genes were detected in *Megasphaera elsdenii*, another species known to use the acrylate pathway for lactate utilization, although the percentage identity was very low for the lactate permease gene (Table S13). Five of the six *lap* cluster genes, were detected in *Anaerotignum propionicum*, excluding *lapB*. This supports the prediction of *lapB* as a lactoyl-CoA epimerase, as *Anaerotignum propionicum* cannot utilize L-lactate [58].

In most butyrate-producing *Firmicutes* species from the human colon, most or all of the genes encoding the central butyrate pathway are present in a single gene cluster, although the butyryl acetate:CoA transferase is separately encoded (Louis & Flint 2009). This was found to be true for *A. soehngenii* but not for *C. catus* GD/7, where the key functions are encoded in at least three separate locations in the genome (Table S14). While the crotonase and beta-hydroxybutyrate dehydrogenase genes are linked in *C. catus* GD/7, two candidate thiolase genes are unlinked. Meanwhile, the butyryl-CoA dehydrogenase (BCD) in the *C. catus* GD/7 genome (GD-7_01569) is situated downstream of the i-LDH (GD-7_01568) and upstream of two ETF genes (GD-7_01570, 01571) (Figure 4A, Table S14). These appear to be homologous with the *lctBCFG* genes of *A. soehngenii*, although these *lct* genes were not upregulated by lactate in *C. catus*.

Of the lactate-utilizing bacteria analyzed, only *Veillonella parvula*, *Bacillus subtilis* and *Campylobacter jejuni* appear to lack the Rnf complex, and all genomes except *C. jejuni* and *Shewanella oneidensis* encoded genes indicative of butyrate and/or propionate production (Figure 5). For bacteria that did not encode L-iLDHs, the lack of a lactate racemase corresponded with an inability to utilize L-lactate. As described earlier, the *lct* and *lap* clusters use different mechanisms of lactate utilization and are present in a different range of organisms. Additionally, *Shewanella oneidensis* uses a third mechanism of lactate utilization, involving a very different lactate dehydrogenase [59]. Homologs of this three-component L-lactate hydrogenase are present in the genomes of *B. subtilis*, *C. jejuni*, *E. coli* and *V. parvula* (Figure 5). Other types of NAD-independent L- and D-lactate dehydrogenases have also been found in organisms from all three domains of life [60]. Thus, there appear to be several divergent paradigms of lactate utilization in bacteria. However, all of these mechanisms appear to require members of the lactate permease superfamily and most possess some member of the acyl-CoA dehydrogenase superfamily. Therefore, the diversity of these proteins warranted detailed further investigation.

**Figure 5.**
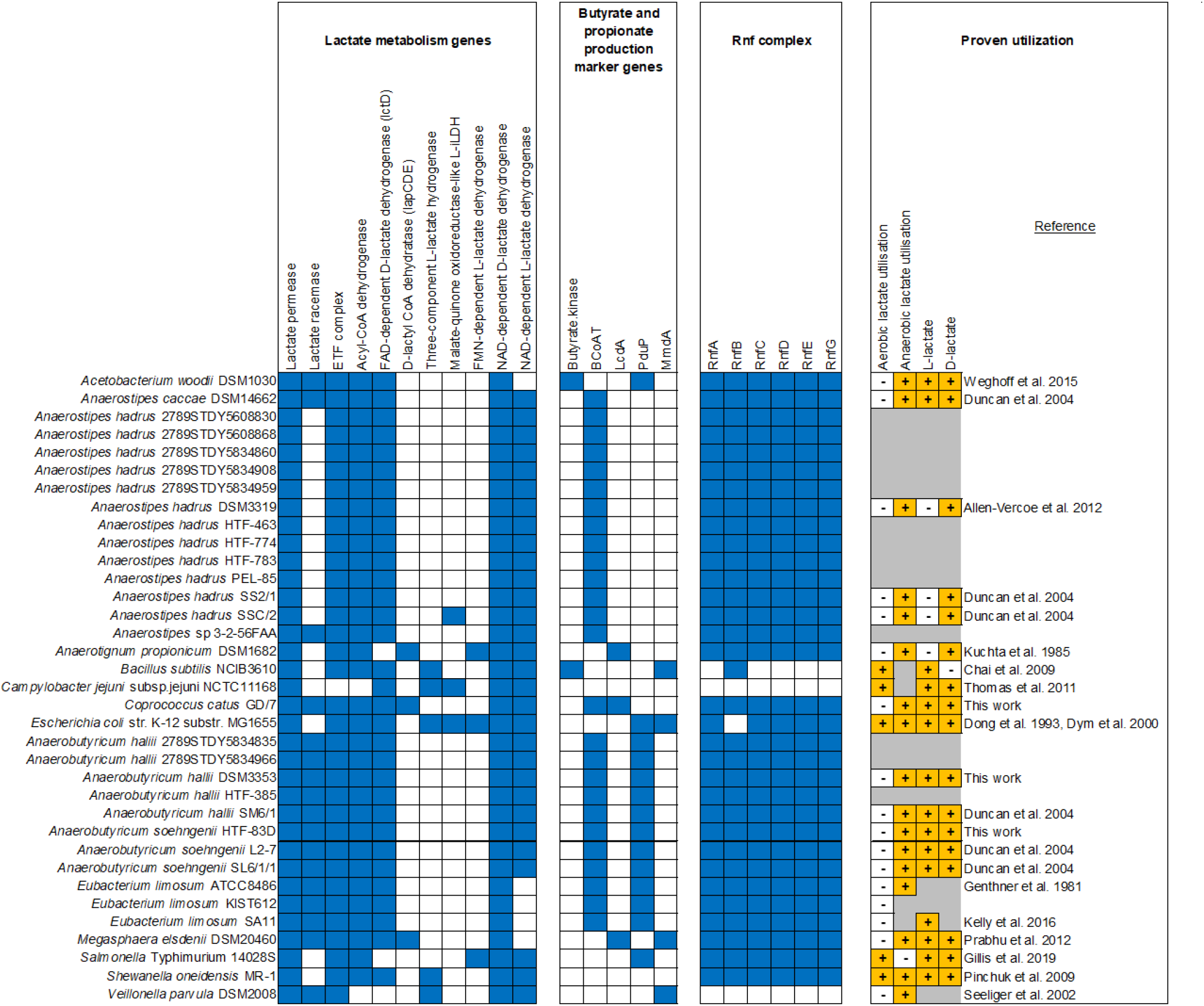
Distribution of lactate utilization and associated genes in lactate-utilizing bacteria. Presence (blue) and absence (white) of genes was determined as described in the Methods section. The ability (+) or inabilty (-) of selected strains to utilize different isomers of lactate during aerobic or anaerobic growth was obtained from the literature [11, 20–22, 58, 59, 61–68]. Certain details could not be recovered for some strains (marked in grey). Full names of genes indicated by short-hand gene nomenclature are as follows: Butyryl-CoA:acetate CoA-transferase (BCoAT), lactoyl-CoA dehydratase, alpha subunit (LcdA), propanediol utilization CoA-dependent propionaldehyde dehydrogenase (PduP), methyl-malonyl-CoA decarboxylase, alpha-subunit (MmdA) and Rnf complex subunits (RnfA-G).

### Phylogeny of lactate permease superfamilies

Phylogenetic analysis of 17653 lactate permeases from bacteria, archaea and eukaryotes revealed five families of lactate permease: LP-I and LP-II, which were present exclusively in bacteria; LP-III and LP-IV, which were present in bacteria and archaea; and LP-V which was present exclusively in eukaryotes. The lactate permeases identified in the transcriptomics work presented here belong to LP-IV, which was further divided into five subfamilies (A-E). Three of the four lactate permease genes in the *C. catus* GD/7 genome, including the copy upregulated on lactate, clustered with the *Anaerotignum* and *S. oneidensis* lactate permeases in LP-IV-C (Figure 6). The fourth clustered with the lactate-induced *A. soehngenii* lactate permease in LP-IV-E. This subfamily notably also contains the lactate permeases of *Anaerostipes* spp., *Megasphaera* spp., *Veillonella* spp., *Acetobacterium* spp. and *E. limosum*.

**Figure 6.**
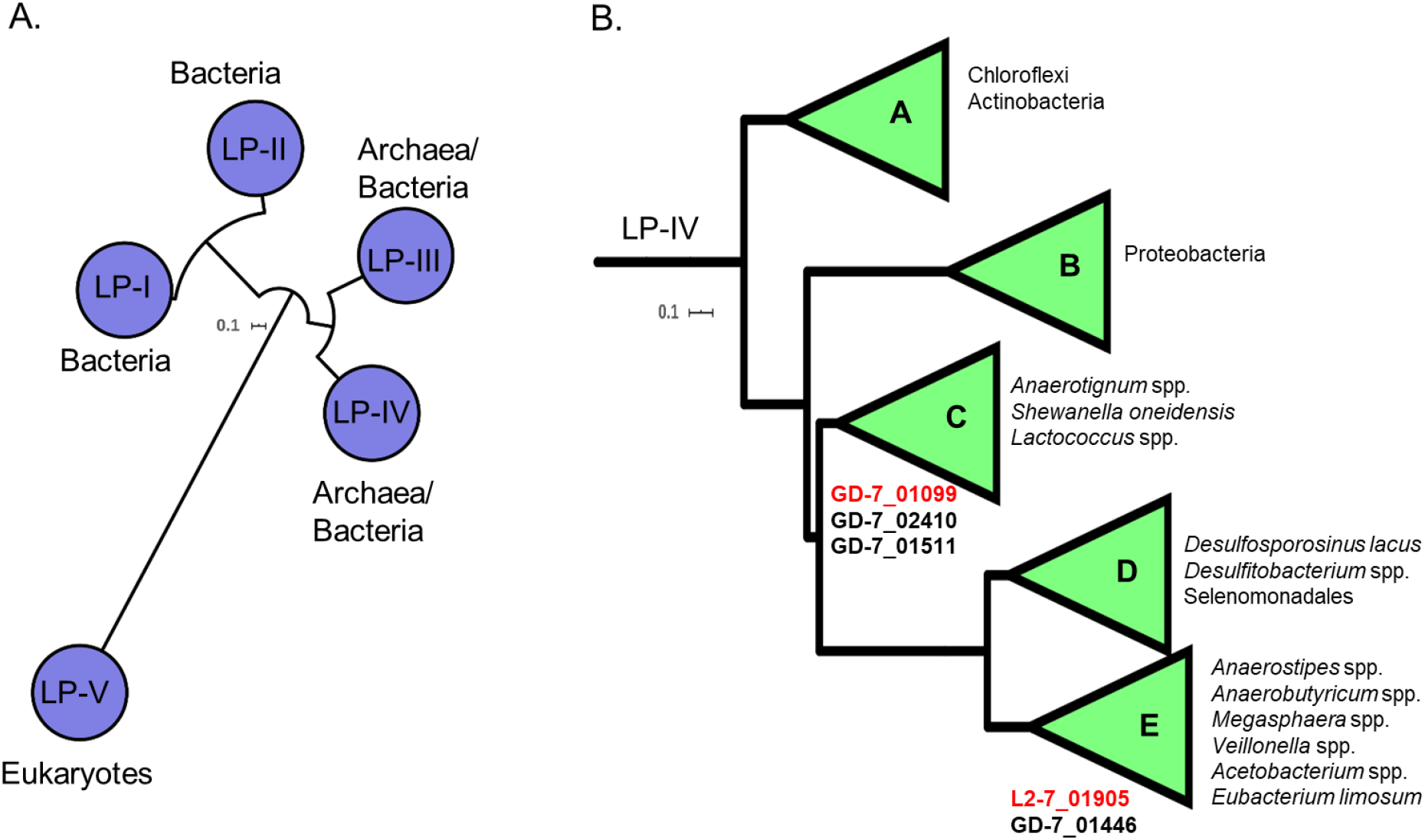
Phylogeny of lactate permease families. A) Families of lactate permease, and B) Sub-families of LP-IV. The conserved lactate permease domain PF02652 was detected in 17653 protein sequences from Uniprot (June 2020). These proteins were formed into 1627 clusters of 70% identity. A maximum-likelihood tree was constructed from the representative sequences of these clusters using the LG+F+R8 model. All branches shown have >70% support of 1000 SH-alrt replicates. Proteins were clustered into protein families and sub-families using the average branch length distances to the leaves in the phylogenetic tree. The tree was rooted using minimal ancestor deviation. Lactate permeases of *C. catus* GD/7 and *A. soehngenii* L2-7 are listed underneath the subfamilies of which they belong to. The upregulated lactate permeases detected in the preceding transcriptomics work in this study are indicated in red font. Representative genes of each family of lactate permease and each sub-family of LP-IV are listed in Table S15 and Table S16, respectively.

Lactate permeases are also notably absent from the genomes of the gut bacteria, *Eubacterium rectale* A1-86, *Roseburia intestinalis* L1-82, *Roseburia hominis* A2-183, *Coprococcus eutactus* L2-50 and *Faecalibacterium praunsnitzii* A2-165, which do not utilize lactate [21], thus indicating a role for lactate permease as a marker gene of lactate utilization, despite disparate mechanisms of lactate oxidation.

### Phylogeny of Acyl-CoA dehydrogenase superfamilies

The acyl-CoA dehydrogenase in the *lct* cluster (L2-7_01909) and its homologs in the *C. catus* and *A. hadrus* genomes were used to create a HMM profile to detect divergent members of this protein superfamily in the genomes of other lactate-utilizing bacteria. These genes were found to be ubiquitous in lactate-utilizing bacteria, with many genomes possessing more than one copy (Table 3). Analysis of these genes revealed that they can be divided into a multitude of clades (Figure 7), many of which, such as the experimentally proven [69] caffeyl-CoA reductase of *A. woodii,* are the only representatives of their clade in our dataset. One clade contained the butyryl-CoA dehydrogenase present in the gene cluster responsible for butyrate production from carbohydrates [70]. This clade also contained the lactate induced *lct* cluster acyl-CoA dehydrogenase, indicating that it too is a butyryl-CoA dehydrogenase. The highly lactate-induced acyl-CoA dehydrogenases of *C. catus* (GD-7_01551 and GD-7_01087) form separate clades (acyl-CoA dehydrogenase families 2 and 4, respectively) with genes from several other species. No genes of these clades have been experimentally characterized, complicating functional prediction. Additionally, genes from *C. catus*, *A. soehngenii* and *A. caccae* form a clade with the experimentally proven [71] acryloyl reductase of *Anaerotignum propionicum*.

**Figure 7.**
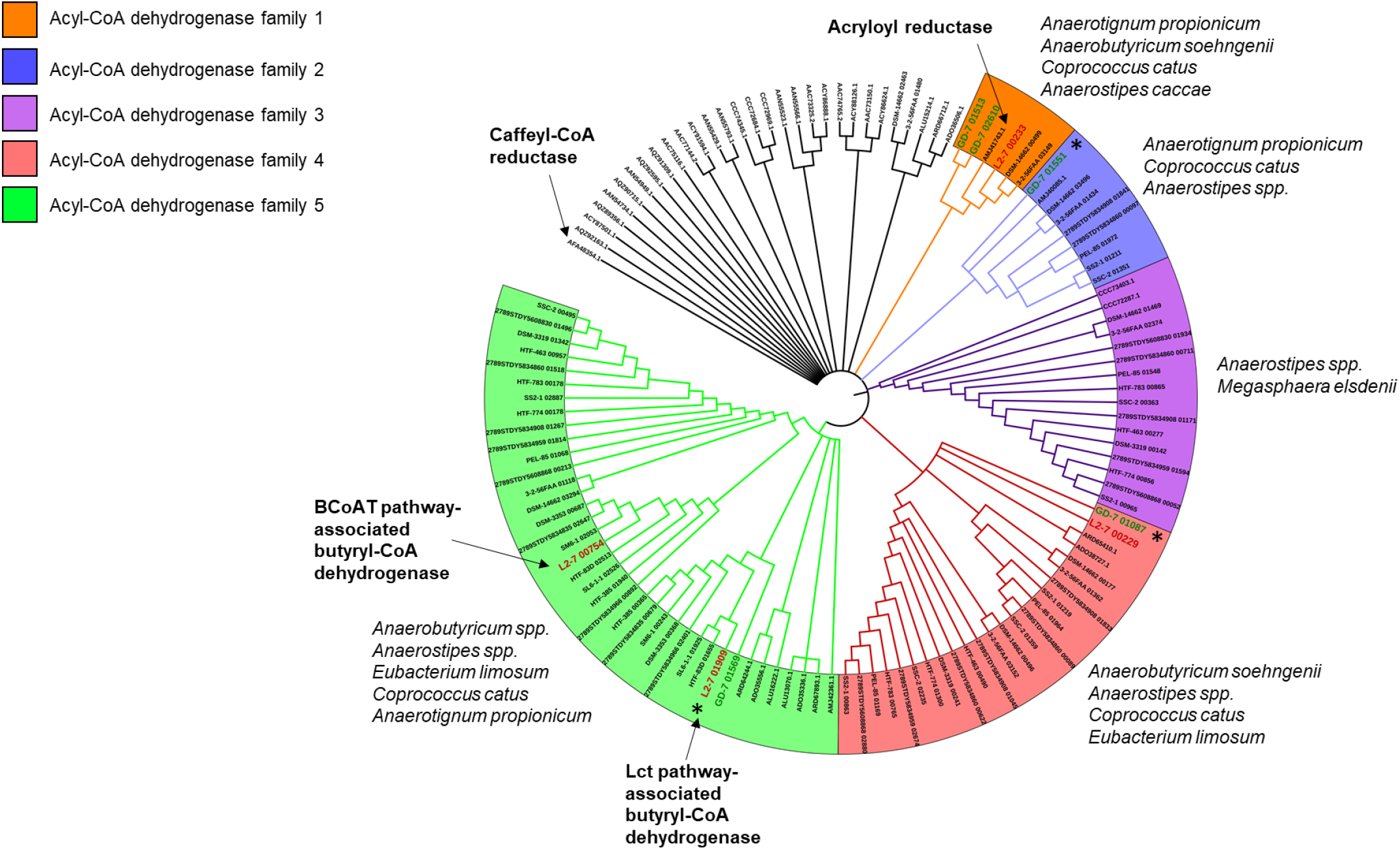
Phylogeny of Acyl-CoA dehydrogenase genes present in lactate-utilizing bacteria. Branch validation was performed using 1000 ultrafast bootstrap replicates and a hill-climbing nearest neighbor interchange (NNI) search was performed to reduce the risk of overestimating branch supports. Branches with less than 90% UFBoot support were collapsed. Genes labelled “Caffeyl-CoA reductase” and “Acryloyl reductase” were experimentally verified by Bertsch *et al.* [69] and Hetzel *et al.* [71], respectively, and the gene labelled “Butyryl-CoA dehydrogenase” is present in the butyrate production loci as described previously [70]. This maximum-likelihood tree was reconstructed using the LG+G4 model. *Anaerobutyricum soehngenii* L2-7 and *Coprococcus catus* GD/7 genes are represented in red and green font, respectively. Asterisks indicate genes that were highly upregulated by lactate in the transcriptomics work presented in this study.

**Table 3.** Acyl CoA dehydrogenase families of lactate-utilizing bacteria. These enzymes are ubiquitous in lactate-utilizing bacteria, with some genomes possessing multiple genes. Many of these genes form five distinct gene families (Figure 7). The number genes of a given family that were upregulated during growth on lactate in *A. soehngenii* L2-7 and *C. catus* GD/7 is shown in parentheses. Black slashes indicate differences in gene copy number between queried genomes of the same species. Expanded to include more lactate-utilizing bacteria genomes in Table S17.

| Acyl-CoA dehydrogenase families |  |  |  |  |  |  |
| --- | --- | --- | --- | --- | --- | --- |
| Species | 1 | 2 | 3 | 4 | 5 | # Genomes queried |
| *Coprococcus catus* | 2 | 1 (1) | 0 | 1 | 1 (1) | 1 |
| *Anaerobutyricum soehngenii* | 0/1 | 0 | 0 | 0/1 | 2 (1) | 3 |
| *Anaerobutyricum hallii* | 0 | 0 | 0 | 0 | 2 | 5 |
| *Anaerostipes caccae* | 1 | 1 | 1 | 2 | 1 | 1 |
| *Anaerostipes* sp. 3-2-56FAA | 1 | 1 | 1 | 2 | 1 | 1 |
| *Anaerostipes hadrus* | 0 | 0/1 | 1 | 0/1/2 | 1 | 12 |
| *Eubacterium limosum* | 0 | 0 | 0 | 0/1 | 0/2 | 3 |
| *Megasphaera elsdenii* | 0 | 0 | 2 | 0 | 0 | 1 |
| *Escherichia coli* | 0 | 0 | 0 | 1 | 0 | 1 |
| *Anaerotignum propionicum* | 1 | 1 | 0 | 0 | 1 | 1 |

*A. soehgnenii* L2-7 and *C. catus* GD/7 possess four and five acyl-CoA dehydrogenases, respectively (Figure 7, Table 3). Two of the *A. soehngenii* L2-7 gene products group with butyryl-CoA dehydrogenase (BCD) enzymes, including the BCD in the central butyrate pathway (L2-7_00754) and the upregulated *lctG* product (L2-7_01919). This strongly suggests that the lactate-induced *lctG* gene product is an alternative BCD enzyme that links reduction of crotonyl-CoA to the oxidation of lactate when lactate is being utilized.

## Discussion

Lactate can be utilized by many aerotolerant bacteria through its conversion to pyruvate via D- or L- lactate oxidases [60, 72]. This reaction is not available to obligate anaerobes however, and different mechanisms are therefore required in the absence of oxygen. As noted previously, anaerobic conversion of lactate to pyruvate is energetically unfavourable [22] and this ability appears to have a limited phylogenetic distribution. Here we have identified two different six-gene clusters (*lct* and *lap*) whose transcriptional expression is highly upregulated during growth on lactate as the sole added carbon and energy source. These two gene clusters correspond to two entirely different mechanisms for lactate utilization. The upregulated cluster in *A. soehngenii* corresponds to five genes of the *lct* gene cluster identified previously in *Acetobacterium woodii*, which is an acetogen that has been shown to convert lactate to pyruvate via a mechanism involving electron confurcation [22]. Products of this *lct* gene cluster were recently detected by proteomic analysis [26], and the cluster reported in the related lactate-utilizing species *A. hallii, A. rhamnosivorans* and *Anaerostipes caccae.* Five of the six clustered genes are also present in *Anaerostipes hadrus,* but all strains of this species with sequenced genomes examined here lacked the lactate racemase (*lctF*), thus explaining why they can only utilize D-lactate.

Our analysis confirms that the *lct* gene cluster is present in the newly sequenced strains of *A. soehngenii*, *A. hallii* and *A. hadrus* described here. Notably, however, the *lct* cluster in all these strains encodes an acyl-CoA dehydrogenase specified by the *lctG* gene that is lacking from the *A. woodii* cluster (Schoelmerich, Katsyv et al. 2018). Since the *lctG* gene product is related to, but distinct from, the BCD gene encoded by the central butyrate pathway cluster [70], it seems likely that it is an alternative butyryl-CoA dehydrogenase (BCD) that acts to reduce crotonyl-CoA when lactate is the growth substrate (Figure 8). While the conversion of pyruvate to acetyl-CoA can produce enough reduced ferredoxin to balance the oxidation of lactate via the D-iLDH/ETF complex [73], redox balance and net energy production are also linked to the operation of the butyrate cycle (Figure 8). The upregulation by lactate of an orthologous enzyme catalyzing the BCD reaction, encoded by *lctG*, may therefore play a critical role. The exact relationships between the co-ordinately upregulated i-LDH, BCD and ETF proteins are not known for *A soehngenii*, although it has been suggested that in *C. butyricum* the LDH-ETF and BCD-ETF complexes act separately [74].

**Figure 8.**
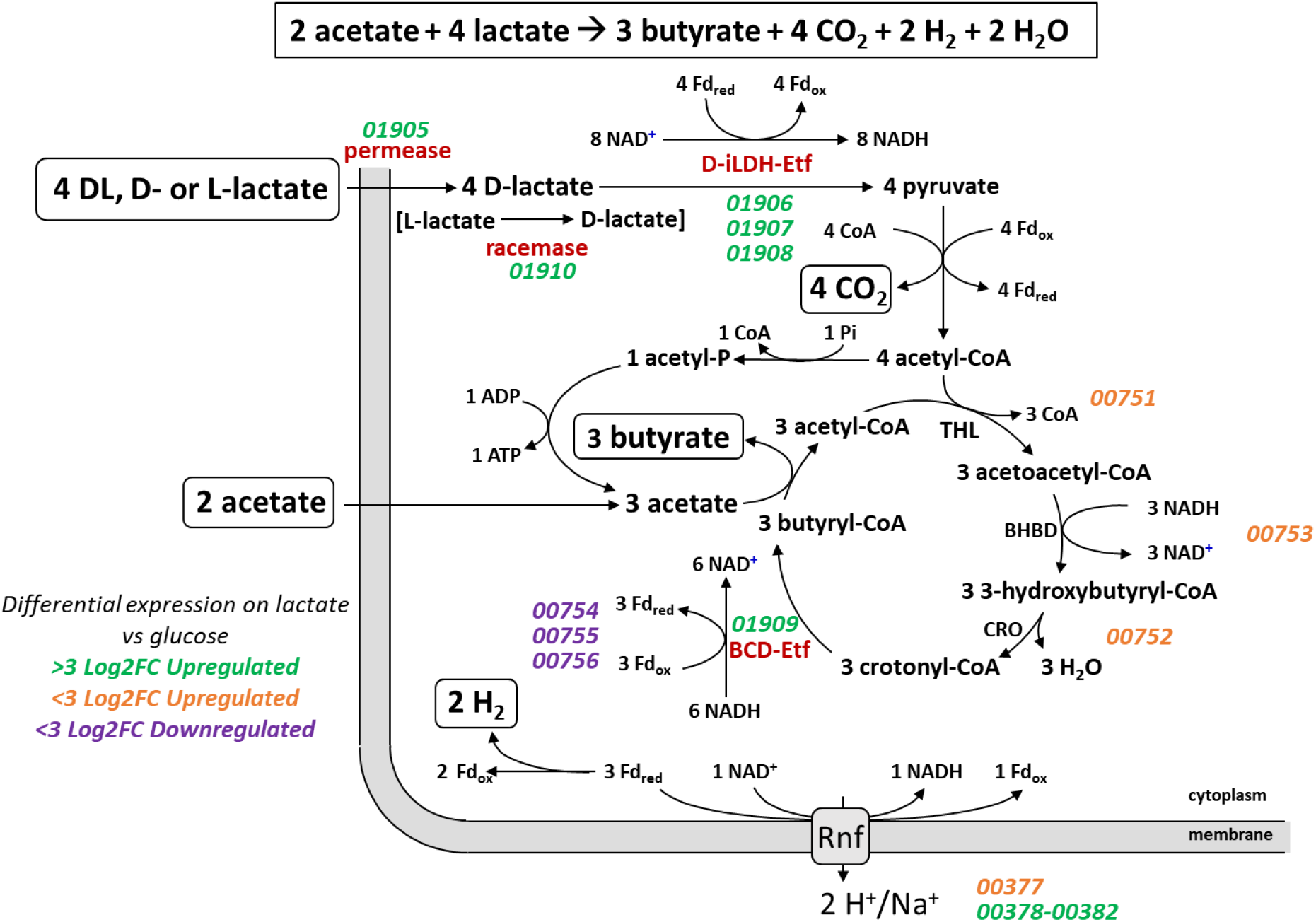
Proposed mechanisms of butyrate formation from lactate in *A. soehngenii* L2-7. Thiolase (THL), β-hydroxybutyryl-CoA dehydrogenase (BHBD) and crotonase (CRO), NAD-independent D-lactate dehydrogenase (D-iLDH), electron transfer flavoprotein (ETF), ferredoxin (Fd) and butyryl-CoA dehydrogenase (BCD). Locus tags are in italics and are colored based on log2 fold change of their expression during growth on lactate versus glucose.

It has been suggested that the molecular hydrogen producing-Hnd complex is involved in balancing redox states during lactate oxidation in *Desulfovibrio* species [57], and its induction during growth of *A. soehngenii* on lactate indicates a likely mechanism for the lactate oxidation-coupled hydrogen production observed in this species [21]. However, resolution of these mechanisms will require further detailed biochemical investigation of the enzymes’ activity and ETF binding of the LctG protein.

The *lap* cluster was identified in *C. catus* GD/7. This cluster has no equivalent in *A. soehngenii* and codes for enzymes required for the acrylate pathway for the conversion of lactate to propionate, notably the three subunits of lactoyl-CoA dehydratase, propionyl-CoA transferase and a presumptive lactoyl-CoA epimerase (Figure 9). Although lactate permeases are present in both the *lap* and *lct* clusters, their sequences are only distantly related to each other. In addition, it seems likely that one of the two unlinked acyl-CoA dehydrogenases that were also upregulated by lactate must correspond to propionyl-CoA dehydrogenase that, along with ETF proteins, carries out the acryloyl-CoA reductase reaction [71].

**Figure 9.**
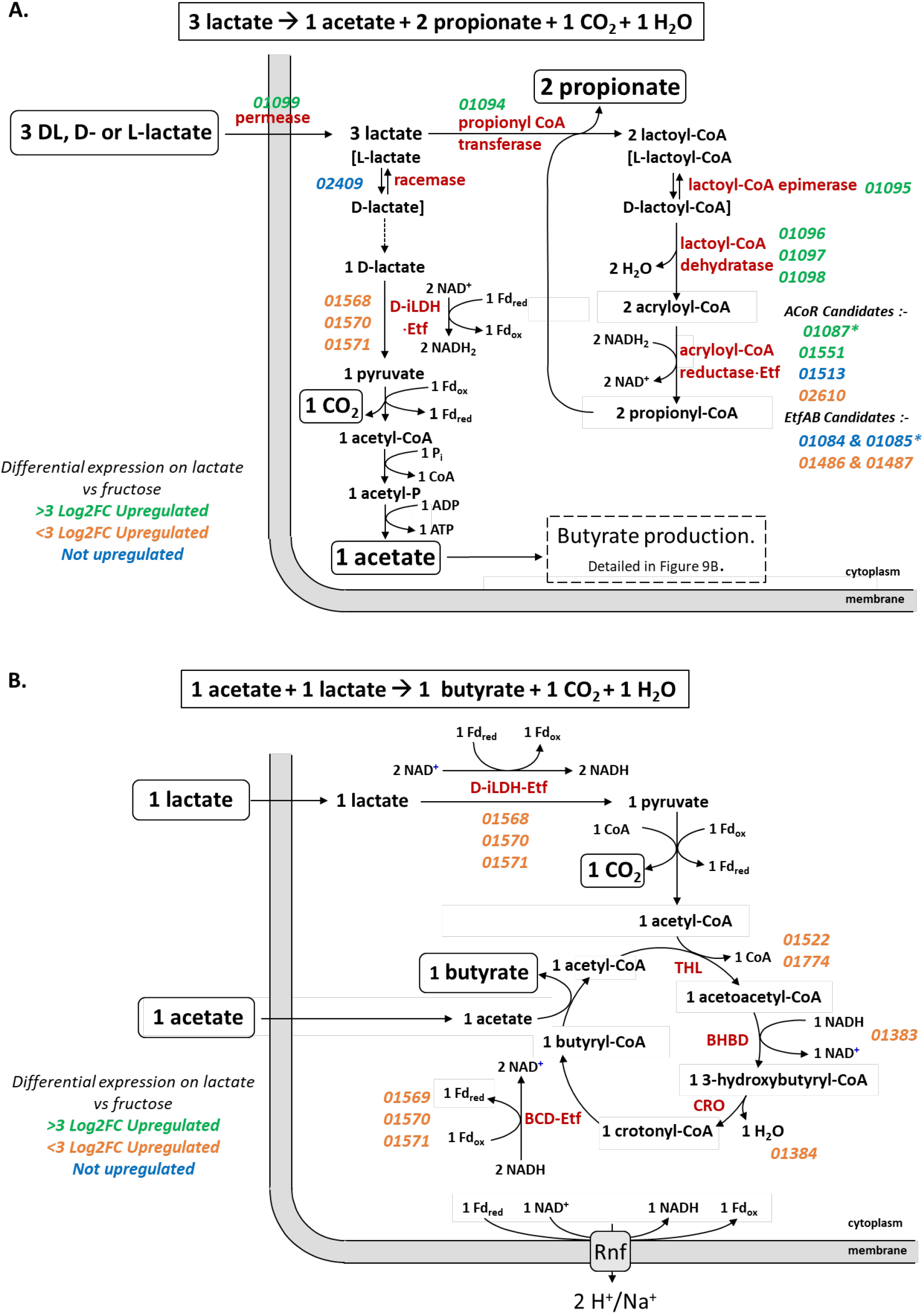
Proposed mechanisms of (A) propionate and acetate formation from lactate and (B) butyrate formation from acetate and lactate in *C. catus* GD/7. NAD-independent D-lactate dehydrogenase (D-iLDH), electron transfer flavoprotein (ETF), ferredoxin (Fd), thiolase (THL), β-hydroxybutyryl-CoA dehydrogenase (BHBD) and crotonase (CRO) and butyryl-CoA dehydrogenase (BCD). Locus tags are in italics and are colored based on log2 fold change of their expression during growth on lactate versus fructose. Asterisks indicate most likely candidate genes.

Operation of the acrylate pathway is normally assumed to result in the conversion of three mol lactate to two mol propionate and one mol of acetate (Figure 9A) [2]. The acetate arises through conversion of lactate to pyruvate via i-LDH, with the reducing equivalents balancing those needed to reduce two acrylate to two propionate. Our results however indicate the formation of some butyrate, in addition to propionate, during growth on lactate. This can be explained readily if we assume that some of the acetate produced is converted along with additional lactate to butyrate (Figure 9B), as happens in *A. soehngenii*. The finding that the i-LDH gene is linked to BCD and ETF genes in *C. catus* GD/7 appears consistent with this proposal. An unexplained feature is the 3-OH butyryl-CoA dehydratase transcript (GD-7_00065) that was upregulated by lactate, although this might possibly be an ortholog of the second β-hydroxybutyryl-CoA dehydrogenase gene (GD-7_01384) that is linked to crotonase in *C. catus* GD/7.

Homologs of *lap* genes were detected in a few other bacterial species, but notably were present in *Megasphaera elsdenii,* which is also known to use the acrylate pathway for lactate utilization. Indeed, *M. elsdenii* is known to convert lactate to butyrate under conditions of carbon-limited growth [66]. It is not known whether the same applies in *C. catus. M. elsdenii* also encodes activities involved in the formation of propionyl-CoA from pyruvate via malate and succinate [75].

Two other anaerobic lactate-utilizing species, *Veillonella parvula* and *Anaerotignum propionicum,* showed high levels of sequence identity with several genes of the *lct* cluster, but lacked a close homolog of the D-iLDH. The latter species however also carries homologs for most of the *lap* cluster and utilizes lactate via the acrylate pathway [76], whereas *V. parvula* encodes a three component L-lactate dehydrogenase, homologous to that of *S. oneidensis* (Figure 5), and uses the ‘randomizing’ succinate pathway rather than the acrylate pathway in producing propionate [2].

In conclusion, we have shown that two distinct routes for anaerobic lactate fermentation in obligate anaerobes, via pyruvate to butyrate and via acrylate to propionate, both involve major transcriptional upregulation of the key genes involved (the *lct* cluster in *A. soehngenii* and the *lap* cluster in *C. catus*) in response to lactate. There was however a clear difference in the impact of hexose sugars on gene expression. Whereas *A soehningii lct* expression was subject to partial repression by glucose, this was not the case for *C. catus* acrylate genes with fructose.

The lack of repression of lactate utilization genes by hexose in *C. catus* may give this and other species with similar metabolism, such as *Megasphaera elsdenii,* particularly important roles in controlling lactate concentrations within gut communities under conditions where there are significant concentrations of free carbohydrates that could be used as alternative energy sources. Indeed, these two species appear to be implicated in accounting for differences in animal productivity that are associated with lactate utilization in the ruminants [14]. The contribution of *Anaerobutyricum* species to lactate utilization in the human colon, where they are relatively abundant within the microbiota, is more likely to be influenced by the availability of sugars as alternative sources of energy. These considerations may be of relevance when selecting strains or strain combinations of lactate-utilizing bacteria intended for use as therapeutic probiotics with the aim of enhancing the stability of the microbial community.

## Supporting information

Collection of all supplementary tables

## Author statements

### Authors and contributors

POS carried out laboratory and *in silico* experiments/data analysis. PL, SHD, HJF and AWW also contributed to experimental design, data interpretation and visualization. ET and HJM generated essential strain data. SS contributed to formal analysis and visualization. POS, PL, SHD, HJF and AWW wrote the manuscript. All co-authors commented on advanced drafts of the manuscript, and contributed edits to the final draft.

## Acknowledgements

The authors would like to thank Donna Henderson (Rowett Institute, University of Aberdeen) for carrying out gas chromatography analysis of short chain fatty acids, and also thank the Centre for Genome-Enabled Biology and Medicine (CGEBM) at the University of Aberdeen for carrying out the Illumina-based transcriptomics sequencing. They would also like to acknowledge the support of the Maxwell computer cluster funded by the University of Aberdeen.

## Supplementary information

**Table S1.** Classifying new *Anaerobutyricum* genomes by average nucleotide identity and average amino acid identity.

**Table S2.** Genome information of strains included in this study. Includes genomes completeness, CDS number and references.

**Table S3.** Short chain fatty acid profiles following growth of lactate-utilizing bacteria on different carbon sources (numerical data for experiments presented in Figure 1).

**Table S4.** Overview showing number of differentially expressed genes during the transcriptomics experiments in each growth condition.

**Table S5.** Differentially expressed genes in *Anaerobutyricum soehngenii* L2-7 during growth on glucose versus lactate

**Table S6.** Differentially expressed genes in *Anaerobutyricum soehngenii* L2-7 during growth on glucose versus glucose and lactate

**Table S7.** Differentially expressed genes in *Anaerobutyricum soehngenii* L2-7 during growth on lactate versus glucose and lactate

**Table S8.** Differentially expressed genes in *Coprococcus catus* GD/7 during growth on fructose versus lactate

**Table S9.** Differentially expressed genes in *Coprococcus catus* GD/7 during growth on fructose versus fructose and lactate

**Table S10.** Differentially expressed genes in *Coprococcus catus* GD/7 during growth on lactate versus fructose and lactate

**Table S11.** Lactate utilization HMM (full sequence bit score >=80) in *Coprococcus catus* GD/7 and *Anaerobutyricum soehngenii* L2-7

**Table S12.** Comparison of *Anaerobutyricum soehngenii* L2-7 lactate utilization locus to that of other known lactate utilizers

**Table S13.** Comparison of *Coprococcus catus* GD/7 lactate utilization locus to that of other known lactate utilizers

**Table S14.** Comparison of *Coprococcus catus* GD/7 butyrate production loci to those of other known lactate utilizers

**Table S15.** Representative genes present in each lactate permease protein family

**Table S16.** Representative genes present in each sub-family of LP-IV

**Table S17.** Abundance of acyl-CoA dehydrogenase families 1-5 in known lactate utilizers

**Figure S1.** Transcriptomic experimental workflow

