## Supplementary figures and images for "Distribution, organization and expression of genes concerned with anaerobic lactate-utilization in human intestinal bacteria"

### Figure S1

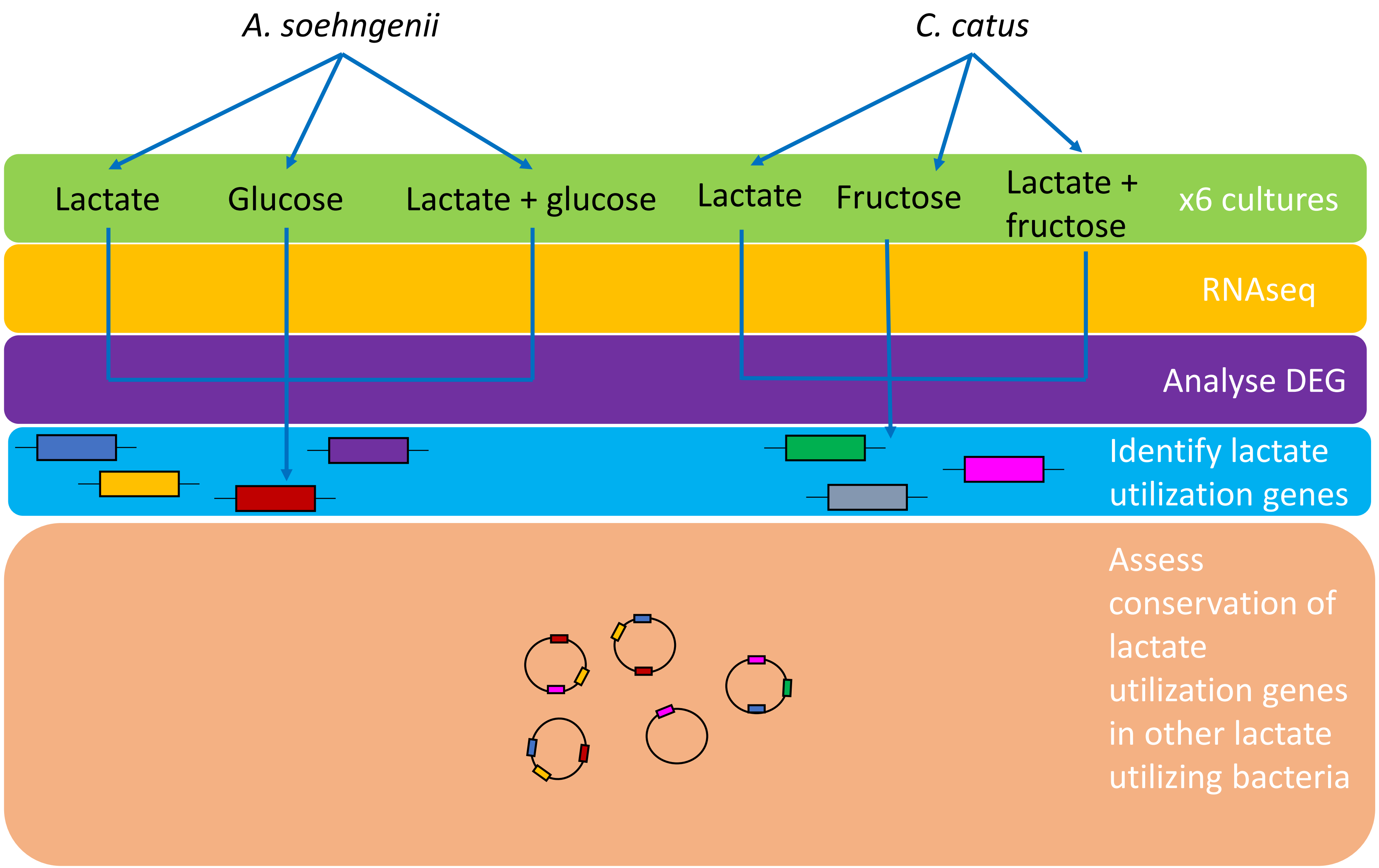
